# FOXC1 and FOXC2 maintain mitral valve endothelial cell junctions, extracellular matrix, and lymphatic vessels to prevent myxomatous degeneration

**DOI:** 10.1101/2023.08.30.555455

**Authors:** Can Tan, Shreya Kurup, Yaryna Dyakiv, Tsutomu Kume

**Affiliations:** Feinberg Cardiovascular and Renal Research Institute, Department of Medicine, Feinberg School of Medicine, Northwestern University, Chicago, Illinois, USA; Honors College, University of Illinois at Chicago, Chicago, Illinois, USA

**Keywords:** mitral valve, FOXC1, FOXC2, endothelial cell-cell junctions, extracellular matrix, lymphatics

## Abstract

**Background:** Mitral valve (MV) disease including myxomatous degeneration is the most common form of valvular heart disease with an age-dependent frequency. Genetic evidence indicates mutations of the transcription factor *FOXC1* are associated with MV defects, including mitral valve regurgitation. In this study, we sought to determine whether murine *Foxc1* and its closely related factor, *Foxc2*, are required in valvular endothelial cells (VECs) for the maintenance of MV leaflets, including VEC junctions and the stratified trilaminar extracellular matrix (ECM).

**Methods:** Adult mice carrying tamoxifen-inducible, endothelial cell (EC)-specific, compound *Foxc1;Foxc2* mutations (i.e., EC-*Foxc*-DKO mice) were used to study the function of *Foxc1* and *Foxc2* in the maintenance of mitral valves. The EC-mutations of *Foxc1/c2* were induced at 7 – 8 weeks of age by tamoxifen treatment, and abnormalities in the MVs of EC-*Foxc*-DKO mice were assessed via whole-mount immunostaining, immunohistochemistry, and Movat pentachrome/Masson’s Trichrome staining.

**Results:** EC-deletions of *Foxc1* and *Foxc2* in mice resulted in abnormally extended and thicker mitral valves by causing defects in regulation of ECM organization with increased proteoglycan and decreased collagen. Notably, reticular adherens junctions were found in VECs of control MV leaflets, and these reticular structures were severely disrupted in EC-*Foxc1/c2* mutant mice. PROX1, a key regulator in a subset of VECs on the fibrosa side of MVs, was downregulated in EC-*Foxc1/c2* mutant VECs. Furthermore, we determined the precise location of lymphatic vessels in murine MVs, and these lymphatic vessels were aberrantly expanded in EC-*Foxc1/c2* mutant mitral valves.

**Conclusions:** Our results indicate that *Foxc1* and *Foxc2* are required for maintaining the integrity of the MV, including VEC junctions, ECM organization, and lymphatic vessels to prevent myxomatous mitral valve degeneration.

## Introduction

Valvular heart disease, which is commonly associated with myxomatous mitral valve degeneration, causes severe regurgitation leading to sudden cardiac arrest or sudden cardiac death. Myxomatous mitral valve degeneration is a non-inflammatory progressive alteration of the mitral valve structure associated with mitral valve prolapse (MVP), a condition characterized by the displacement of one or both leaflets of the mitral valve into the left atrium during the contraction phase of the heart ^1,2^. MVP affects more than 7 million Americans, whereas up to 25% of individuals with MVP will develop degenerative mitral regurgitation. MVP is a heterogeneous disease, and its pathogenesis is not fully understood. Although recent genetic studies have identified causative genes associated with MVP ^3-7^, little is known about the etiology of defects in the mitral valve that lead to the progression of MVP.

Studies using mutant mouse models are critical for elucidating the mechanisms underlying mitral valve development and disease. In mice, mitral valve development initiates immediately after looping the heart tube that consists of myocardial (outer) and endocardial (inner) cell layers ^8^. A subpopulation of endocardial endothelial cells (ECs) undergoes endothelial-to-mesenchymal transition (EndMT) and migrates into the cardiac jelly of the atrioventricular canal. Following EndMT, valve mesenchymal cells (i.e., interstitial cells) and valvular ECs (VECs) proliferate and produce the extracellular matrix (ECM) to promote the elongation and maturation of the valve leaflet primordia until birth. Postnatally, the mitral valve remodels to form three layers of stratified ECM ^8^, an elastin-rich atrialis layer, a proteoglycan-rich spongiosa layer, and a collagen-rich fibrosa layer.

Recent single cell RNA-sequencing (scRNA-seq) studies demonstrated molecularly distinct populations of VECs, interstitial cells (ICs), and other cell types ^9,10^. A specific population of valve ECs on the fibrosa side of the mitral valve leaflet specifically expresses the transcription factor *Prox1* ^9,11,12^, which is a key regulator of lymphatic EC specification/identity and is upregulated by lymph flow ^13^. Importantly, structural abnormalities in the mitral valve are often associated with dysregulated ECM. Although valve ICs are known to produce ECM components, the contribution of VECs to mitral valve diseases such as myxomatous degeneration remains largely unknown. How VECs participate in the maintenance of mitral valve integrity in the adult, including the stratified trilaminar ECM structure, has yet to be elucidated.

FOXC1 and FOXC2 are closely related members of the FOX transcription factor family and have numerous essential roles in cardiovascular development, health, and disease ^14^. Mutations or changes in the copy number of human *FOXC1* are associated with autosomal-dominant Axenfeld-Rieger syndrome, which is characterized by abnormalities in the eye and extraocular defects ^15^, while inactivating mutations of human *FOXC2* are responsible for the autosomal dominant syndrome Lymphedema-distichiasis, which is characterized by obstructed lymph drainage in the limbs and the growth of extra eyelashes ^16^. Importantly, there is genetic evidence that human *FOXC1* mutations are associated with mitral valve defects, including mitral valve regurgitation ^17,18^, whereas a recent study shows that mice with *Foxc2* knockdown in valvular endothelial cells do not exhibit any mitral valve abnormalities ^11^. Although our previous studies show that both *Foxc1* and *Foxc2* are expressed in endocardial ECs in the developing mouse heart ^19-21^, it remains to be elucidated whether *Foxc1* and *Foxc2* deficiency in mitral VECs impairs the maintenance of mitral valve integrity, including VEC junctions and ECM components.

Employing an EC-specific *Foxc1/Foxc2* double mutant (i.e., EC-*Foxc*-DKO) line, we have recently demonstrated that FOXC1 and FOXC2 play cooperative roles in blood and lymphatic ECs in various organs such as the small intestine and mesentery ^22,23^. In this study, adult EC-*Foxc*-DKO mice were used to determine the requirement for *Foxc1* and *Foxc2* in the maintenance of mitral valve leaflets because our initial analyses revealed both genes are highly expressed in VECs of mitral valves in adult mice. We found that adult EC-deletions of *Foxc1* and *Foxc2* lead to dysregulated ECM components, accompanied by abnormal formation of reticular adherens junctions in VECs of mitral valves and reduced expression of PROX1 in a subset of VECs. Moreover, we also determined the distribution of lymphatic vasculature in mitral valves, and EC-*Foxc*-DKO mice exhibited expanded lymphatic vessels in mitral valves.

Taken together, the results from our new studies suggest that lack of *Foxc1* and *Foxc2* in VECs damages the maintenance of mitral valve integrity, leading to impairments in VEC junctions, ECM organization, and lymphatic vessels, therefore causing myxomatous degeneration of mitral valves.

## Materials and Methods

The data that support the findings of this study are available within the article (and in the Supplemental Material).

### Animal husbandry and treatment

*Foxc1*^*fl/fl*^*;Foxc2*^*fl/fl* 24^ and *Cdh5-Cre*^*ERT2* 25^ mice were used. EC-specific compound *Foxc1;Foxc2* mutant mice were generated by crossing *Foxc*-floxed females (*Foxc1*^*fl/fl*^*;Foxc2*^*fl/fl*^) with *Cdh5-Cre*^*ERT2*^*;Foxc1*^*fl/fl*^*;Foxc2*^*fl/fl*^ (EC-*Foxc*-DKO) males as described previously ^22^. For Cre recombination efficiency detection, *mTmG/+;Cdh5-Cre*^*ERT2*^*;Foxc1*^*fl/fl*^*;Foxc2*^*fl/fl*^ (mTmG/EC-*Foxc*-DKO) mice were generated by crossing mTmG females (*mTmG/mTmG;Foxc1*^*fl/fl*^*;Foxc2*^*fl/fl*^) with EC-*Foxc*-DKO males. Genotyping of mice was performed by Transnetyx Inc. To induce gene deletion of *Foxc1* and *Foxc2* in ECs, 7–8-week-old male adult mice were treated with 40 mg/mL tamoxifen (Tm, Cayman Chemical #13258) in corn oil at a dose of 150 mg/kg by oral gavage once daily for 5 consecutive days.

### Whole-mount (WM) staining

Whole-mount staining of mitral valves (MVs) was performed as previously described^23^ with slight modifications. Briefly, after fixation, the dissected MVs together with the connected mitral annulus and the papillary muscle (PM) were permeabilized in PBST (0.3% Triton X-100 in PBS) for 1h at 4°C, blocked in blocking buffer (5% donkey serum, 0.5% BSA, 0.3% Triton X-100, 0.1% NaN_3_ in PBS) for 2 h at 4°C, and then incubated with the indicated primary antibodies (Table S1) diluted in the blocking buffer for 2∼3d at 4°C. Samples were washed with PBST for several times, followed by incubation with indicated fluorochrome-conjugated secondary antibodies (Table S1) diluted in the blocking buffer for 2d at 4°C. The samples were washed again with PBST for several times, post-fixed with 4% PFA. Most of the PM was removed before the MVs were cleared with FocusClear (CelExplorer Labs #FC-101) and flat-mounted on slides in mounting medium.

### Statistics

For quantification, statistical analysis was performed using GraphPad Prism 8.0 (GraphPad Software). *P* values were obtained by performing Mann-Whitney *U* test. Data are presented as box-and-whisker plots of representative experiments from at least three biological replicates. *P* values <0.05 were considered statistically significant.

## Results and Discussion

### Expression of FOXC1 and FOXC2 in the mitral valve of the mouse heart

Previous histological studies demonstrate that murine *Foxc1* is expressed in the mitral valve (MV) of the developing heart ^18,21^. When evaluated via quantitative real-time RT-PCR (qPCR), *Foxc1* was more highly expressed than *Foxc2* in CD45^−^/CD31^+^ cardiac ECs isolated from the adult mouse hearts (Figure 1A). Furthermore, both FOXC1 and FOXC2 proteins were detected in mitral VECs of adult mice with their expression more abundant in VECs on the ventricular (fibrosa) side of the MV leaflets, whereas FOXC1 was also highly expressed in ICs as well (Figure 1B).

**Figure 1.**
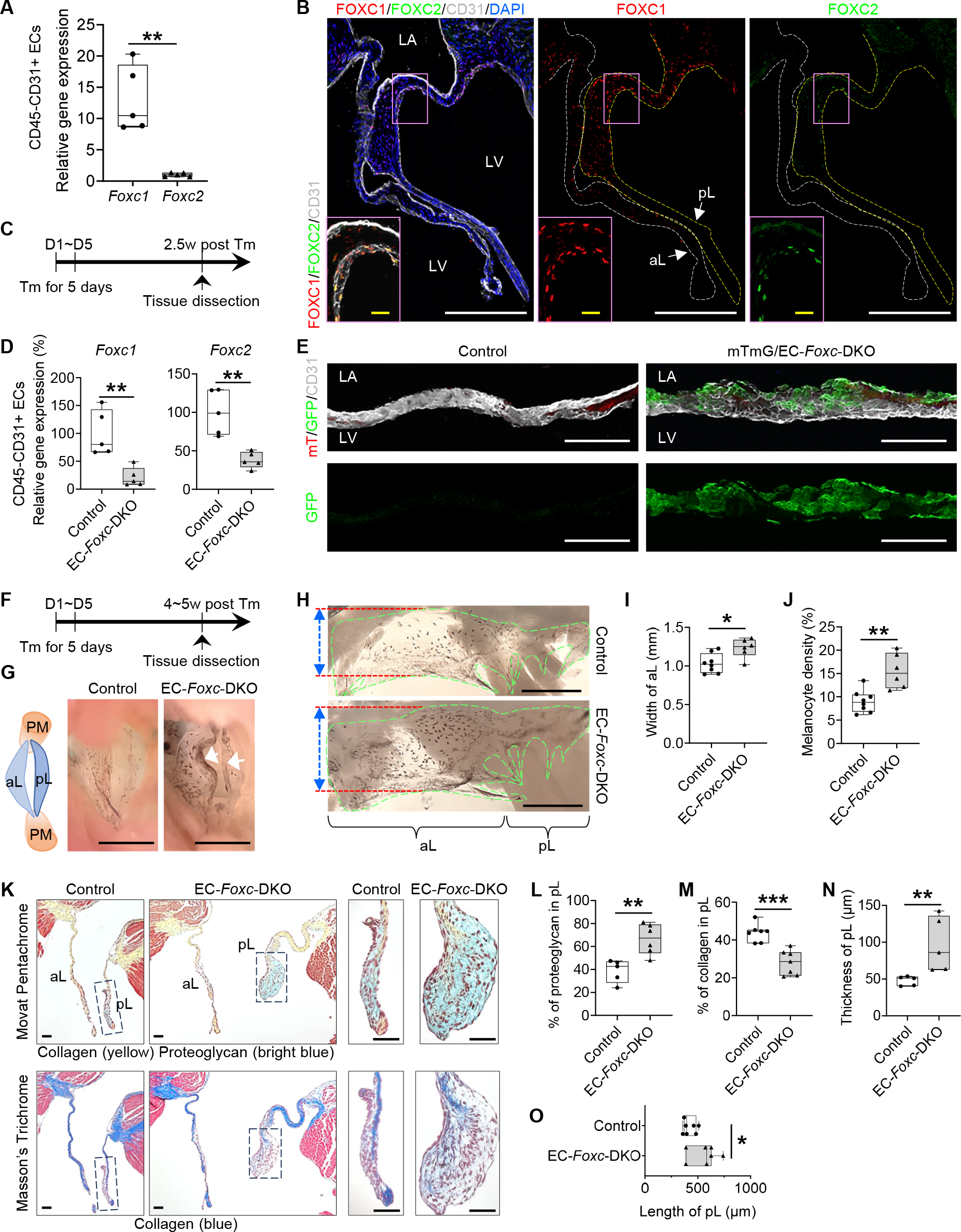
EC-specific deletion of *Foxc1* and *Foxc2* leads to myxomatous degeneration of mitral valves. **(A)** Relative mRNA expression of *Foxc1* and *Foxc2* in isolated CD45^-^CD31^+^ endothelial cells (ECs) from *Foxc1*^*fl/fl*^*;Foxc2*^*fl/fl*^ mouse heart. Data are box-and-whisker plots, Mann– Whitney *U* test, each symbol represents one mouse, N = 5, \*\**P* < 0.01. **(B)** Representative fluorescent images of mitral valves (10 µm of cryo-section) from *Foxc1*^*fl/fl*^*;Foxc2*^*fl/fl*^ mouse stained with CD31 (white), FOXC1 (red) and FOXC2 (green). White and yellow broken lines outline the anterior (aL) and posterior leaflets (pL) of the mitral valve (MV), respectively. In MV, FOXC1/FOXC2 expressing cells are not only ECs located on both sides (FOXC1) or fibrosa side (ventricle-facing side) (FOXC2) of the MV (especially pL), but also a few interstitial cells. LA: left atrium; LV: left ventricle. White/yellow scale bars = 200 or 20 µm, respectively. **(C)** Schematic showing the time of tamoxifen (Tm) injection and tissue dissection for Figure 1, D-E. **(D)** Relative mRNA expression of *Foxc1* and *Foxc2* in isolated CD45^-^CD31^+^ ECs from hearts post Tm dose in mice (Control: *Foxc1*^*fl/fl*^*;Foxc2*^*fl/fl*^, EC-*Foxc*-DKO: *Cdh5-Cre*^*ERT2*^*;Foxc1*^*fl/fl*^*;Foxc2*^*fl/fl*^). Data are box and whisker plots, Mann-Whitney *U* test, each symbol represents one mouse, N = 5, \*\**P*<0.01. **(E)** Cre recombination efficiency detection in mTmG/EC-*Foxc*-DKO (*mTmG/+;Cdh5-Cre*^*ERT2*^*;Foxc1*^*fl/fl*^*;Foxc2*^*fl/fl*^) mice and littermate control mice (*mTmG/+;Foxc1*^*fl/fl*^*;Foxc2*^*fl/fl*^) by immunostaining of frozen heart sections with GFP and CD31. Representative confocal z-stacked images of these thick sections (16 µm) show GFP expression in ECs on MV. Scale bars = 100 µm. **(F)** Schematic showing the time of tamoxifen (Tm) injection and tissue dissection for Figure 1, G-O, and Figure 2. **(G)** Representative images of MVs taken from the perspective of the left ventricle. Arrows show the accumulation of melanocytes at the free edge of the leaflets in EC-*Foxc*-DKO mouse. The structural diagram in the left panel shows the structures seen in the right panels. PM: papillary muscle; aL: anterior leaflet; pL: posterior leaflet. Scale bar = 1 mm. **(H-J)** Representative images **(H)** of flat-mount MVs (atrial aspect) under a stereo microscope. MVs are transparent membranes outlined by green broken lines. The darker background is cardiac muscles at the back of the MVs. The black dots or areas on the MVs are melanocytes. Red lines and double-sided arrows indicate the width of the anterior leaflet (aL) of the MV. pL: posterior leaflet. Scale bars = 1 mm. The width of aL and the density of melanocytes (%=area of melanocytes/area of MVs x 100%) were quantified in **(I)** and **(J)**, respectively. Data are box-and-whisker plots, Mann–Whitney *U* test, each symbol represents one mouse, N = 6∼8, \**P* < 0.05, \*\**P* < 0.01. **(K-O)** Representative images **(K)** of Movat Pentachrome and Masson’s Trichrome stained MVs in serial sections show the ECM components including proteoglycan and collagen in MVs. Scale bars = 50 µm. The percentages of proteoglycan and collagen in posterior leaflets (pLs) were quantified in **(L)** and **(M)**. The thicknesses **(N)** and lengths **(O)** of the pLs were also measured and quantified. Data are box-and-whisker plots, Mann–Whitney *U* test, each symbol represents one mouse, N = 5∼7, \**P* < 0.05, \*\**P* < 0.01, \*\*\**P* < 0.001.

### Generation of tamoxifen-inducible, EC-specific, *Foxc1/c2*-mutant mice

Since we found both FOXC1 and FOXC2 are expressed in the mitral VECs, we analyzed tamoxifen-inducible, EC-specific, compound *Foxc1;Foxc2*-mutant (*Cdh5*-*Cre*^*ERT2*^;*Foxc1*^*fl/fl*^*;Foxc2*^*fl/fl*^) mice, which (after the mutation is induced in the adult) will be referred to as EC-*Foxc*-DKO mice ^22,23^. To induce the mutations, adult (7ꟷ8-week-old) mice were treated with tamoxifen (150 mg/kg) by oral gavage for 5 consecutive days, and 2.5 weeks after tamoxifen treatment (Figure 1C), qPCR analysis confirmed that *Foxc1* and *Foxc2* expression were significantly less common in CD45^−^/CD31^+^ cardiac ECs of EC-*Foxc*-DKO mice than in the corresponding cells of their control littermates (Figure 1D). Since our EC-*Foxc*-DKO mice express Cre from the EC-specific *Cdh5* promoter ^25^, we crossed them with dual Rosa26mTmG reporter mice ^26^; then, we treated their adult offspring with tamoxifen as described above and identified recombined EGFP^+^ VECs in the MVs and confirmed the efficiency of Cre-mediated recombination (Figure 1E). Of note, the MVs of mTmG/EC-*Foxc*-DKO mice were abnormally shaped and much thicker than those of the control mice (see below for more details).

### EC-specific deletion of *Foxc1* and *Foxc2* impairs the structure of mitral valves

We further examined structural changes in the mitral valves in adult EC-*Foxc*-DKO mice compared to their littermate controls 4∼5 weeks after tamoxifen treatment (Figure 1F). Our initial investigation indicated structural abnormalities in EC-*Foxc1/c2* mutant MVs (Figure 1G), and isolated MVs of EC-*Foxc*-DKO mice were much wider and had an increased number of melanocytes, compared to the control mice (Figure 1H through J). Given evidence that the number of melanocytes localized in the murine MV leaflets contributes to the mechanical properties such as stiffness ^27^, we next examined alterations in ECM components of the MVs of EC-*Foxc*-DKO mice via combined analyses of Movat Pentachrome and Masson’s Trichrome staining (Figure 1K). The MVs of EC-*Foxc*-DKO mice had altered ECM components with increased proteoglycan and decreased collagen (Figure 1L and 1M), and they were thicker and longer than the controls (Figure 1N and 1O), which suggested myxomatous degeneration of MVs. Milder structural abnormalities were found in MVs of either EC-*Foxc1* or *Foxc2* single KO mice (data not shown). Additionally, mitral annulus dilation was found in EC-*Foxc*-DKO mice, with increased area of mitral annulus opening and larger distance across the roots of MV leaflets than in control mitral annulus (Figure S1A through S1D), which was a potential cause for mitral regurgitation. Given the interactions between VECs and ICs in mitral valve leaflets ^28^, these results suggest that *Foxc1/c*2 expression in VECs contributes to the maintenance of the mitral valve ECM.

### Disrupted EC junctions in the mitral valves of mice with EC-specific deletion of *Foxc1/c2*

We next examined whether EC-*Foxc1/c2* deletions impair the maintenance of EC junctions in mitral valves because we and others previously found defective lymphatic EC junctions in lymphatic valves of *Foxc1* and/or *Foxc2* mutant mice ^22,29^. Whole mount immunostaining for CD31 and VE-cadherin and confocal imaging first revealed notable protrusions in the MVs of EC-*Foxc*-DKO mice (Figure 2A, arrows), indicative of leaflet thickening and redundancy found in MVP. Remarkably, the size, shape, and arrangement of control CD31+ VECs on both atrial and fibrosa sides of posterior leaflet (pL) differed in a zone-dependent manner (Figure S1E and S1F), and similar EC zones were found in anterior leaflet. In contrast, EC-*Foxc*-DKO mice showed elongated EC Zone 1 (Figure S1G), in which the size and number of *Foxc*-mutant VECs were smaller and increased, respectively (Figure S1H). More significantly, the reticular adherens junctions (AJ) formed at the interface of two overlapping VECs (Figure 2B, arrows), as previously shown in cultured ECs ^30,31^, were found mainly in EC Zone 3 and 4 in both control and EC-*Foxc*-DKO pLs, but appeared in Zone 1 and 5 in pLs of EC-*Foxc*-DKO mice (Figure 2B and Figure S1E). The extent of VECs containing reticular AJs was much increased in EC-*Foxc*-DKO mice (Figure 2C). The presence of the reticular AJs was further confirmed by the orthogonal view of z-stacked images (Figure 2D) as well as 3D-reconstructed images (Figure 2E and 2F). It should be noted that VE-cadherin+ junctions in the reticular AJs of EC-*Foxc*-DKO mice were abnormally discontinued (Figure 2F, arrows), accompanied by reduced VE-cadherin levels (Figure 2G). The formation of reticular AJs in ECs is thought to retain low permeability ^30,31^, while cultured lymphatic ECs under oscillatory shear stress are known to form such cellular structures to stabilize cell-cell junctions ^29^. Thus, it is likely that *Foxc1/c2*-deficient VECs of mitral valves aberrantly form reticular AJs as a compensatory mechanism, but these cell junctions are disrupted, as seen in lymphatic ECs in vivo ^22^.

**Figure 2.**
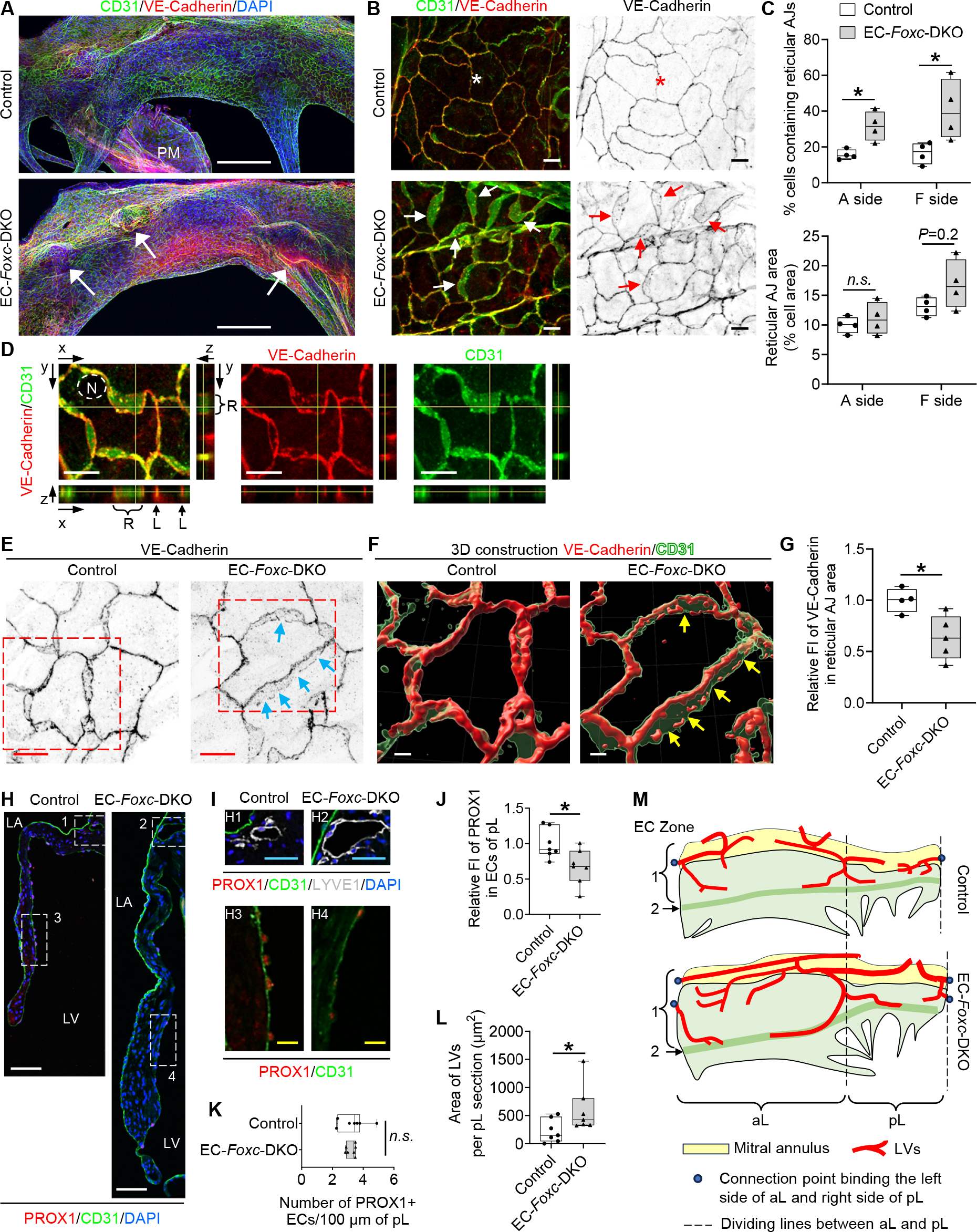
FOXC1 and FOXC2 maintain the integrity of endothelial cell junctions and lymphatic vessels in the mitral valves. **(A)** Representative confocal images of the whole-mount posterior leaflets (pLs) of MVs stained with endothelial cell markers CD31 (green) and VE-Cadherin (red), as well as nuclear dye DAPI (blue). Images of maximum intensity projections of the atrial aspect of pLs show thickened pL with obvious protrusions (indicated by arrows) in EC-*Foxc*-DKO mice. Note that the locally increased VE-Cadherin staining in EC-*Foxc*-DKO pL is due to the images of thick pL being stacked. PM: papillary muscle. Scale bars = 200 µm. **(B)** Magnified confocal images of whole-mount pLs stained with CD31/VE-Cadherin showing cell-cell junctions of ECs on the fibrosa side of pLs (in EC Zone 5 as shown in Supplemental Figure 1F). Stars indicate the linear junctions while arrows indicate the reticular adherens junctions (AJs) of ECs. Scale bars = 10 µm. **(C)** Quantification of the reticular AJs on both atrial (A) and fibrosa (F) sides of the pLs based on the images as shown in Figure 2A. Data are box-and-whisker plots, Mann– Whitney *U* test, each symbol represents one mouse, N = 4, \**P* < 0.05, *n*.*s*., not significant. **(D)** Orthogonal view of z-stacked images (0.3 um/step) showing the cell-cell junctions (stained with VE-Cadherin and CD31) of endocardial ECs in pL of EC-*Foxc*-DKO mouse. Yellow lines on the stack indicate the point in the stack that is being analyzed. R: reticular AJ area; L: linear cell-cell junction. N: nucleus of EC. Scale bars = 10 µm. **(E-G)** Representative confocal images **(E)** of ECs containing reticular AJs stained with VE-Cadherin in pLs of MVs. The confocal images (boxed area) were further reconstructed to 3D isosurfaces by Imaris Workstation as shown in **(F)**. Arrows in E and F indicate the discontinued cell-cell junctions in the reticular AJs. Scale bars = 10 µm in E and 4 µm in F. **(G)** Quantification of fluorescent intensity (FI) of VE-Cadherin in reticular AJs of ECs in pLs based on the images as shown in Figure 2E. Data are box-and-whisker plots, Mann– Whitney *U* test, each symbol represents one mouse, N = 4∼5, \**P* < 0.05. **(H-L)** Representative fluorescent images **(H and I)** of pLs of MVs. Box 1 and 2 in **H** and their magnified images (H1 and H2) in **I** show the lymphatic vessels located at the atrial side of the root of MVs, where the mitral annulus area is also positioned as shown in Figure S2A, stained by lymphatic endothelial cell (LEC) markers PROX1/CD31/LYVE1. Box 3 and 4 in **H**, as well as their magnified images (H3 and H4) in **I**, show the decreased expression of PROX1 in ECs at the fibrosa side of pL in EC-*Foxc*-DKO mouse. LA: left atrium, LV: left ventricle. White/blue/yellow scale bars = 50/25/10 µm, respectively. Quantification was performed based on the fluorescent images as shown in Figure 2H for: relative fluorescent intensity (FI) of PROX1 in ECs on the fibrosa side of pL **(J)** and the number of PROX1^+^ ECs **(K)**, as well as the area of lymphatic vessels (LVs) in pL sections **(L)**. Data are box-and-whisker plots, Mann–Whitney *U* test, each symbol represents one mouse, N = 7, \**P* < 0.05, *n*.*s*., not significant. **(M)** Structural diagram (based on Figure S2D) shows the distribution of lymphatic vessels (LVs) in both anterior (aL) and posterior (pL) leaflets of MVs. In control mouse, LVs start from the blind-ended capillaries underneath the lower part of EC Zone 1. The lymphatic capillaries join at the dividing lines of aL and pL, then enter the mitral annulus to form collecting lymphatic vessels. The mitral annulus is also covered by EC zone 1 cells. In EC-*Foxc*-DKO mouse, most of the blind ends of LVs are underneath EC zone 2.

Furthermore, PROX1 levels in the VECs on the fibrosa side of MVs were significantly reduced in EC-*Foxc*-DKO mice (Figure 2H through 2J) while the number of PROX1+ VECs was not altered (Figure 2K). Since a recent study indicates that *Prox1* acts upstream of *Foxc2* in aortic valves ^11^, the downregulation of *Prox1* in *Foxc1/c2*-mutant VECs may be attributable to a feedback loop.

### Expansion of lymphatic vessels in the mitral valves of EC-*Foxc1/c2* mutant mice

Limited evidence indicates that lymphatic vessels (LVs) are present in murine MVs ^32^, while other mammalian species such as humans, pigs, and dogs have lymphatic vasculature in the MVs ^33,34^. We therefore examined the distribution of LVs in MVs of control and EC-*Foxc*-DKO mice via co-immunostaining of the lymphatic markers, PROX1 and LYVE1 (Figure 2I) in paraffin sections. Careful examination revealed that in both control and EC-*Foxc*-DKO mice, most of the LVs were located at the atrial side of the root of MV leaflets (Figure 2H and 2I), where the mitral annulus area was also positioned (Figure S2A). Importantly, consistent with our previous finding that *Foxc1* and *Foxc2* deletion causes enhanced lymphangiogenesis ^35^, EC-*Foxc*-DKO mice had dilated and higher numbers of LVs (Figure 2I and 2L; Figure S2B). Notably, additional LVs were found in the middle of the pLs (underneath EC Zone 2) of EC-*Foxc*-DKO mice (Figure S2C). Whole mount immunostaining further confirmed the distribution of LVs in MVs (Figure S2D and Figure 2M). The lymphatic capillaries extended to EC Zone 2 in MVs of EC-*Foxc*-DKO mice whereas the control mice had LVs only in EC Zone 1.

Collectively, we show here that *Foxc1* and *Foxc2* are essential for the integrity of MV leaflets by maintaining VEC junctions, ECM organization, and LVs (Figure S3). To our knowledge, this is the *first* study to demonstrate reticular AJs in MVs in vivo. Our study also reveals the precise location of LVs in murine MVs, which are expanded in EC-specific mutant mice for *Foxc1* and *Foxc2*.

## Acknowledgments

We thank William Muller (Northwestern University) for helpful advice. *Cdh5-Cre*^*ERT2*^ mice were provided by Dr. Ralf Adams at the Max-Planck-Institute for Molecular Biomedicine, Germany. Confocal imaging work was performed at the Northwestern University Center for Advanced Microscopy supported by NCI CCSG P30 CA060553 awarded to the Robert H Lurie Comprehensive Cancer Center. This study was supported by National Institutes of Health (NIH) grants RO1HL144129, R01HL159976, and R01EY034740 (to T. Kume).

## Supplemental Material

Figure S1-S3

Supplemental Methods^23^

Table S1

