## Supplemental Material for "FOXC1 and FOXC2 maintain mitral valve endothelial cell junctions, extracellular matrix, and lymphatic vessels to prevent myxomatous degeneration"

### Supplemental Figures

#### Figure S1

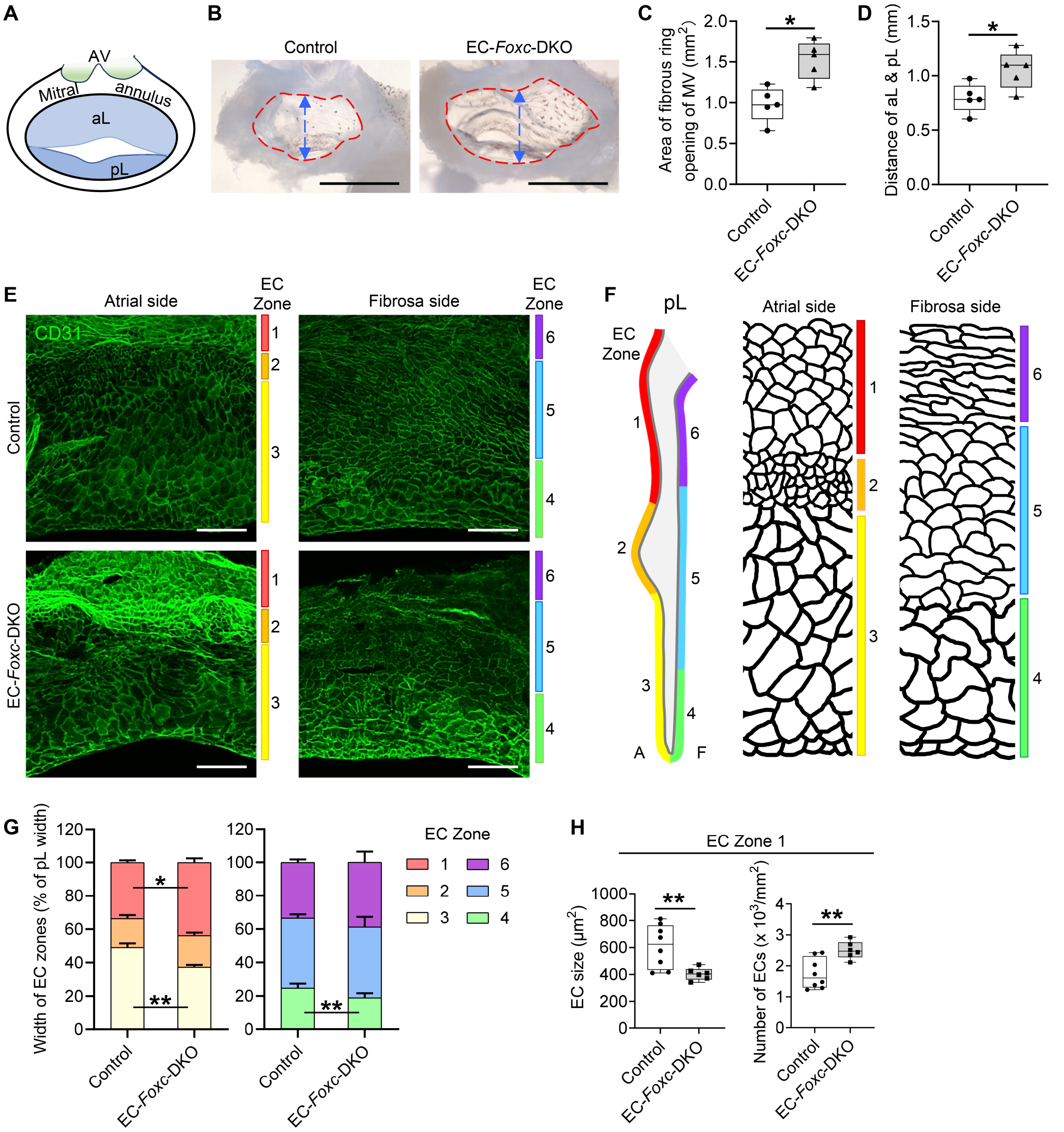

**Figure S1. FOXC1 and FOXC2 are required for the arrangement of ECs in the mitral valves.**

**(A-D)** Structural diagram **(A)** showing the anterior (aL) and posterior leaflets (pL) as well as the mitral annulus (fibrous ring) and aortic valve (AV) presented in B. **(B)** Representative images of mitral annulus of MVs taken from the perspective of atrium under stereo microscope, showing enlargement of the mitral annulus opening in EC-*Foxc*-DKO mice 4~5 weeks post Tm treatment. Scale bars = 1 mm. The area of mitral annulus opening (indicated by red broken circles) and the distance across the roots of aL and pL (indicated by blue double-sided arrows) were measured and quantified in **(C)** and **(D)**, respectively. (C) and (D), Data are box and-whisker plots, Mann-Whitney *U* test, each symbol represents one mouse, N = 5, *$P$<0.05.

**(E)** Representative confocal z-stacked images of whole-mount posterior leaflets (pLs) of MVs stained with CD31 show the arrangement of ECs at both atrial and fibrosa sides of pLs in mice at 4~5 weeks post Tm treatment. The ECs on the pL were divided into 6 different zones according to their size, shape and arrangement as illustrated in F. The ECs were disorganized in MVs of EC-*Foxc*-DKO mouse. Scale bars = 100 µm.

**(F)** Illustration of EC Zones at both atrial (A) and fibrosa (F) sides of posterior leaflet (pL) of MV in control mouse. Left panel: ECs are arranged from Zone 1-3 at the atrial side, to Zone 4-6 at fibrosa side. EC Zone 1 and Zone 6 are close to the root of pL, while Zone 3 and Zone 4 are close to the free edge of pL. ECs are in small size in Zone 2, medium size in Zone 1, 5 and 6, large size in Zone 3 and 4. ECs are polygon in Zone 1-5 but adopt an elongated shape along the mitral annulus in Zone 6. The reticular AJs can be found mostly in Zone 3 and 4, while linear cell-cell junctions are mostly distributed in Zone 1, 2, 5 and 6.

**(G)** Quantification of width of different EC zones in pLs based on the confocal images as shown in E. Data are Mean ± SEM in a stacked percentage bar chart, unpaired *t* test, N = 4 and 5 in control and EC-*Foxc*-DKO group, respectively, *$P$<0.05, **$P$<0.01.

**(H)** Quantification of size and number of ECs in Zone 1. Data are box-and-whisker plots, Mann–Whitney *U* test, each symbol represents one mouse, N = 6~8, *$P$ < 0.05, **$P$ < 0.01.

#### Figure S2

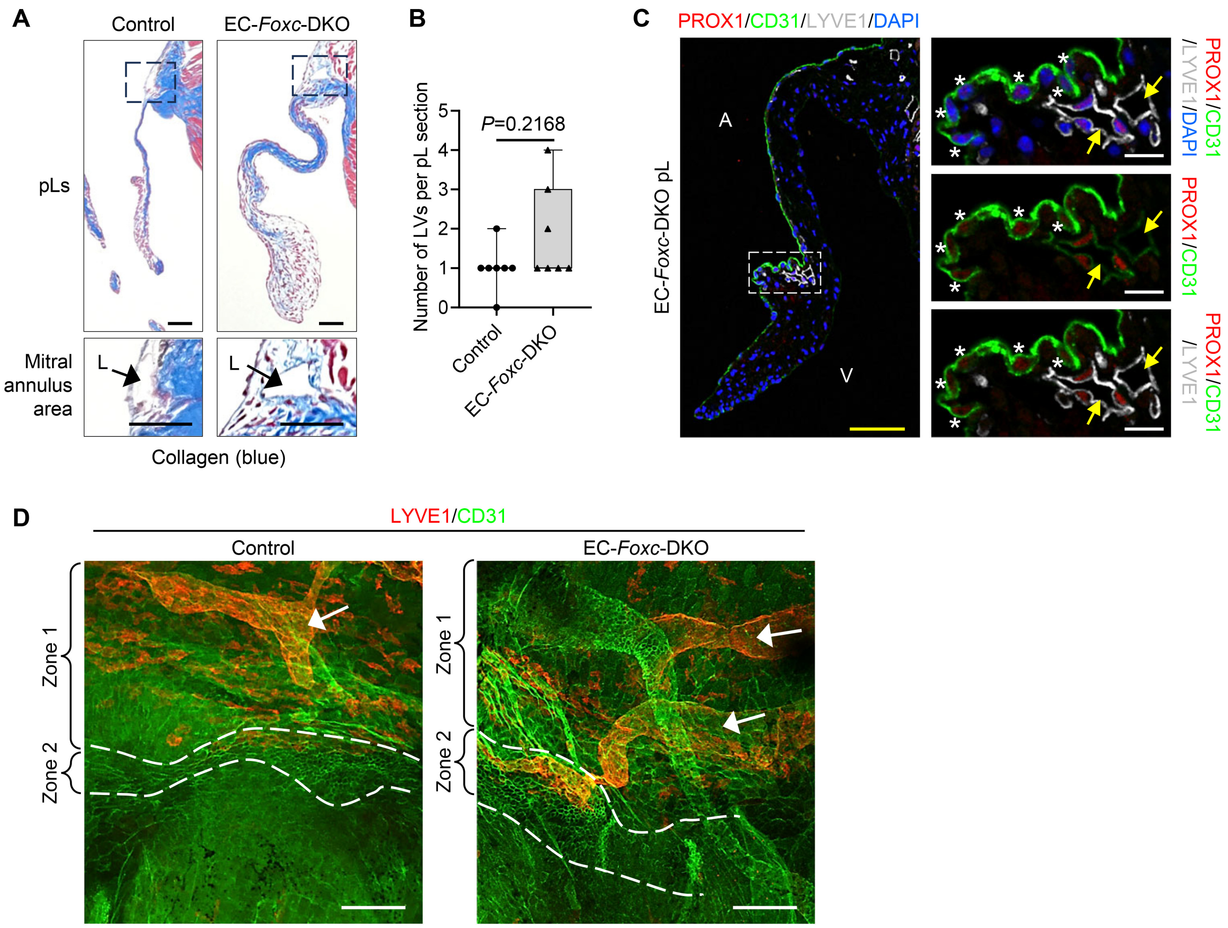

**Figure S2. FOXC1 and FOXC2 are required for the maintenance of lymphatic vessels in the mitral valves.**

**(A)** Representative images of Masson's Trichrome staining of posterior leaflets (pLs) of MVs. Magnified area focuses on the mitral annulus area (as shown in Supplemental Figure 1A) which is at the root of the pL and rich in collagen. Lymphatic vessels (L, as shown in Figure 2, H and I) were found in the mitral annulus area. Scale bars = 50 μm.

**(B)** Quantification of number of lymphatic vessels (LVs) based on the fluorescent images as shown in Figure 2H. Data are box-and-whisker plots, Mann–Whitney *U* test, each symbol represents one mouse, N = 7.

**(C)** Representative fluorescent images of LVs (arrows) found in the center part of pL (underneath EC Zone 2) in EC-*Foxc*-DKO mouse. Lymphatic endothelial cells (LECs) were stained with their markers PROX1/CD31/LYVE1. Stars indicate the PROX1$^+$ ECs found in EC Zone 2 only at the atrial side of pL. Yellow/white scale bars = 50 or 10 μm, respectively.

**(D)** Representative confocal z-stacked images of whole-mount pLs stained with LYVE1 (red) and CD31 (green) show the distribution of lymphatic vessels (LVs indicated by arrows). Increased number of LVs and dilated LVs were found in EC-*Foxc*-DKO mice. In control mice, most of the LVs were located underneath EC Zone 1; while the LVs extend to EC Zone 2 (zone with small size of ECs) in EC-*Foxc*-DKO mice. The broken lines outline the EC Zone 1 and 2 according to the criteria described in Supplemental Figure 1F. Scale bars = 100 μm.

#### Figure S3

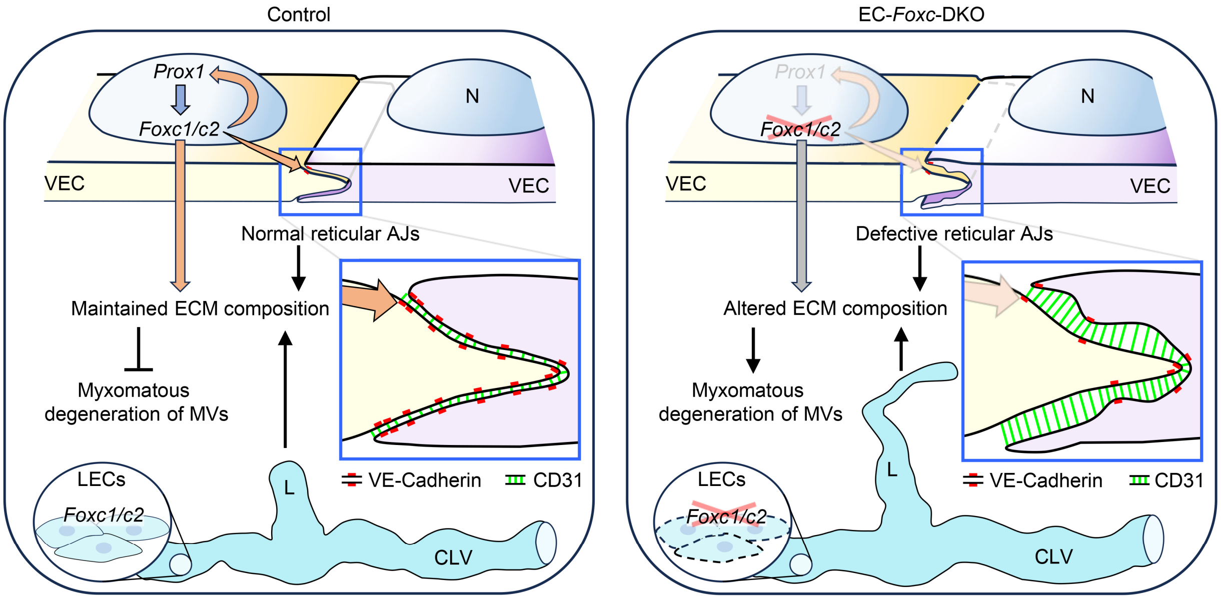

**Figure S3. *Foxc1* and *Foxc2* are required in VECs and LECs to prevent myxomatous degeneration of mitral valves.**

Schematic drawing of the mechanism by which *Foxc1/c2* maintain the cell-cell junctions of valvular endothelial cells (VECs), extracellular matrix (ECM) composition and lymphatic vessels to prevent myxomatous degeneration of mitral valves (MVs). In normal (control) VECs, *Foxc1/c2* are regulated by *Prox1*, which can be conversely regulated by *Foxc1/c2* to maintain ECM composition via cell cross-talks between VECs and valvular interstitial cells. Meanwhile, *Foxc1/c2* regulate the expression of VE-Cadherin to maintain normal reticular adherens junctions (AJs) of VECs, which may also help to maintain the ECM composition to prevent myxomatous degeneration of MVs. *Foxc1/Foxc2* expression in lymphatic endothelial cells (LECs) maintain the structure and function of collecting lymphatic vessels (CLVs) and the blind-ended lymphatic capillaries (L), which is also important to maintain the ECM composition. In EC-*Foxc*-DKO mice, knock out of *Foxc1/c2* in VECs leads to the defective reticular AJs in VECs and altered ECM composition. Deletion of *Foxc1/c2* in LECs leads to the dilation of CLVs and elongated lymphatic capillaries, which affects the lymphatic function and causes altered ECM composition, hence resulting in myxomatous degeneration of MVs. N: nucleus of VEC.

#### Supplemental Methods

##### Cre recombination efficiency detection

To detect Cre recombination efficiency, mTmG/EC-*Foxc*-DKO mice and their littermate control Cre negative mTmG mice were used. 2.5 weeks after Tm treatment, the hearts were harvested and fixed in 4% paraformaldehyde (PFA), followed by dehydration in 30% sucrose and OCT (Sakura Finetek, USA) embedding. 16 μm cryosections were cut and immunostained with CD31 and GFP antibodies (Table S1) and a nuclear-specific dye DAPI. EGFP fluorescent signal was detected by imaging to evaluate the Cre recombination efficiency.

##### Tissue collection

Four to five weeks post Tm treatment, the hearts were collected for histological analysis. Briefly, transcardial perfusion was performed on the adult mice with cold PBS followed by 4% PFA after anesthesia. The hearts were then dissected and post-fixed in 4% PFA at 4°C for 4 h (for frozen or whole-mount samples) or for overnight (O/N, for paraffin embedded samples). The fixed hearts were then processed to OCT- or paraffin-embedded samples. For whole-mount staining of mitral valves (MVs), after the fixation of heart, the MVs (anterior and posterior leaflets) together with their connected mitral annulus and papillary muscles (PM) were dissected and processed to the staining.

##### Movat Pentachrome staining and Masson's Trichrome staining

Movat Pentachrome and Masson's Trichrome staining were performed on 4-μm paraffin sections of the hearts according to the manufactures' instructions (Scytek Laboratories, MPS2; and Thermo Scientific, #87019).

**Immunohistochemistry (IHC) staining**

IHC-P on 4 μm paraffin (P) sections and IHC-F on 10~16 μm frozen (F) sections were performed as previously described[23]. The antibodies used are listed in Table S1.

**Imaging**

To image the mitral annulus with MV leaflets, the fixed and dissected MVs together with the mitral annulus and PM structures were embedded in 1% agarose (Sigma, A9414-10G) in PBS. Images were taken from the perspective of atrium under stereo microscope (Nikon SMZ 745) connected to a camera from Nikon DS-Fi2 (for quantification) or to the camera of Apple iPhone 12 Pro Max (for representative images). Flat-mount mitral valve images were obtained using the same imaging system. Movat Pentachrome and Masson's Trichrome staining images were acquired using an Olympus Vanox AHBT3 Research Microscope (Tokyo, Japan). WM and IHC staining images were acquired using a Nikon A1 Confocal Laser Microscope with the software of NIS-Elements Viewer. Images were processed and analyzed with Adobe Photoshop, Fiji (ImageJ) and Imaris Workstation (for 3-D reconstruction).

**Endothelial cell isolation from mouse heart**

Endothelial cells were isolated from mouse heart for further qPCR analysis as previously described[23] with slight modifications. Briefly, the heart was collected after cardiac perfusion with cold PBS. The heart was washed in DMEM and processed upon dissociating into single cell suspension in a digestion buffer (1.5 mg/mL Collagenase type I, 60 U/mL DNase I, 60 U/mL hyaluronidase, 100 units/mL penicillin and 100 μg/mL streptomycin in DMEM) for 30 min at 37°C, followed by filtration through a 70 μm and a

40 µm cell strainer. After several washes, the cell suspension was then incubated with Dynabeads labeled with CD45 antibody (Biolegend #103102) to deplete the $CD45^+$ cell population. The $CD45^-$ cell suspension was then incubated with Dynabeads labeled with CD31 antibody (BD #553369) to get the $CD45^-CD31^+$ ECs. Finally, the sorted ECs were used for RNA isolation and qPCR analysis.

**RNA isolation and qPCR analysis**

The RNeasy Mini Kit (Qiagen #74104) was used for RNA extraction from cells. The concentration of RNA was determined using NanoDrop™ 2000 Spectrophotometers (Thermo Scientific). cDNA was synthesized using an iScript reverse transcriptase kit (Bio-Rad #170-8891). qPCR was performed on triplicates of cDNA samples by using QuantStudio® 3 Real-Time PCR System (Applied Biosystems), Fast SYBR reaction mix (Applied Biosystems), and gene specific primer sets. 18S was used as internal standards for mRNA expression in mouse samples. *Foxc1* primer sequences are forward: 5’-TTCTTGCGTTCAGAGACTCG-3’, and reverse: 5’-TCTTACAGGTGAGAGGCAAGG-3’. *Foxc2* primer sequences are forward: 5’- AAAGCGCCCCTCTCTCAG-3’, and reverse: 5’-TCAAACTGAGCTGCGGATAA-3’. *18S* primer sequences are forward: 5’- GAAACTGCG AATGGCTCATTAAA-3’, and reverse: 5’- CCACAGTTATCCAAGTAGGAGAGGA-3’.

**Quantification**

Fiji (ImageJ) software was used for the measurement of length, area, number of cells and fluorescent intensity (FI) of specific markers.

The width of anterior leaflet (aL) of MVs was determined by examining the flat-mount MV images and by measuring the vertical distance between the root of aL and its free edge as shown in Figure 1H. The length of posterior leaflet (pL) of MVs was determined by examination of serial sections and by measuring the distance along the center of the leaflet from the valvular root to its tip. The thickness of pLs was measured at the thickest point perpendicular to the length axis of the leaflet.

For the quantification of percentages of proteoglycan and collagen in pL, only one color was applied to the heart paraffin sections: bright blue (Alcian Blue solution, pH 2.5) from Movat Pentachrome staining for proteoglycan, or blue (Anline Blue solution) from Masson's Trichrome staining for collagen, respectively. Images were taken for serial sections focusing on the leaflet area. Area of the single color for proteoglycan (bright blue) or collagen (blue) was measured by thresholding in Fiji. The area of pL was also measured by selecting the region of pL. The percentage of proteoglycan or collagen in pL was then calculated: % = area of proteoglycan or collagen / area of pL x 100%.

For the quantification of percentage of cells containing reticular AJs, about 300~800 total cells were counting and examined for a single side (atrial or fibrosa) of the pL, based on the confocal z-staked images taken under a 20x objective. The area of reticular AJs (% cell area) was also quantified: % = area of reticular AJs / area of cells containing reticular AJs x 100%.

For quantification of FI of VE-Cadherin (in reticular AJs) and PROX1 (in VECs on fibrosa side of pL), about 15~20 reticular AJs (for VE-Cadherin) and 25~35 VECs (for PROX1) were quantified respectively for each sample based on the immunostaining images taken under a 20x objective.

**Study approval**

All experimental protocols and procedures used in this study were approved by the Institutional Animal Care and Use Committee (IACUC) at Northwestern University.

#### Table S1

| Primary Antibodies | | | | | |
|---|---|---|---|---|---|
| **Antibody** | **Supplier** | **Catalog number** | **Host Species** | **Clonality** | **Application** |
| CD31 | BD Biosciences | 553370 | Rat | mAb | WM, IHC-F |
| CD31 | Cell signaling | 77699 | Rabbit | mAb | IHC-P |
| CD31 | R&D | AF3628 | Goat | pAb | WM |
| FOXC1 | Abcam | ab227977 | Rabbit | mAb | IHC-F |
| FOXC2 | Kind gift from Dr. N Miura (Miura et al., 1997, Genomics) | - | Rat | mAb | IHC-F |
| GFP | Abcam | ab13970 | Chicken | pAb | IHC-F |
| LYVE-1 | Abcam | ab14917 | Rabbit | pAb | WM |
| LYVE-1 | eBioscience | 14-0443-82 | Rat | mAb | IHC-P |
| PROX1 | R&D | AF2727 | Goat | pAb | IHC-P |
| VE-Cadherin | BD | 555289 | Rat | mAb | WM |
| VE-Cadherin | R&D | AF1002 | Goat | pAb | WM |

| Secondary Antibodies | | | | |
|---|---|---|---|---|
| **Antibody** | **Reactivity** | **Host Species** | **Supplier** | **Application** |
| Alexa 488-conjugated | Goat | Donkey | Thermo Fisher | WM, IHC-P, IHC-F |
| Alexa 488-conjugated | Rat | Donkey | Thermo Fisher | |
| Alexa 488-conjugated | Chicken | Donkey | Thermo Fisher | |
| Alexa 488-conjugated | Rabbit | Donkey | Thermo Fisher | |
| Alexa 568-conjugated | Rabbit | Donkey | Thermo Fisher | |
| Alexa 568-conjugated | Goat | Donkey | Thermo Fisher | |
| Alexa 568-conjugated | Rat | Donkey | Abcam | |
| Alexa 647-conjugated | Rat | Donkey | Thermo Fisher | |
| IHC-P: immunohistochemistry staining on paraffin sections; IHC-F: Immunohistochemistry staining on frozen sections; WM: Whole-mount staining. | | | | |

**Table S1. Antibodies used for whole-mount and sections**
